# Metagenomics for chronic meningitis: clarifying interpretation and diagnosis

**DOI:** 10.1101/213561

**Authors:** Michael R. Wilson, Brian D. O’Donovan, Jeffrey M. Gelfand, Hannah A. Sample, Felicia C. Chow, John P. Betjemann, Maulik P. Shah, Megan B. Richie, Mark P. Gorman, Rula A. Hajj-Ali, Leonard H. Calabrese, Kelsey C. Zorn, John E. Greenlee, Jonathan H. Blum, Gary Green, Lillian M. Khan, Debarko Banerji, Charles Langelier, Chloe Bryson-Cahn, Whitney Harrington, Jairam R. Lingappa, Niraj M. Shanbhag, Ari J. Green, Bruce J. Brew, Ariane Soldatos, Luke Strnad, Sarah B. Doernberg, Cheryl A. Jay, Vanja Douglas, S. Andrew Josephson, Joseph L. DeRisi

**Author notes:** Authors contributed equally (Michael R. Wilson, Brian D. O’Donovan, Jeffrey M. Gelfand). Corresponding Author: Michael Wilson, UCSF Weill Institute for Neurosciences, Department of Neurology, 675 Nelson Rising Lane, NS212A, Campus Box 3206, San Francisco, CA.

## Abstract

**Importance:** Identifying infectious causes of subacute and chronic meningitis can be challenging. Enhanced, unbiased diagnostic approaches are needed.

**Objective:** To present a case series of patients with diagnostically challenging subacute and chronic meningitis in whom metagenomic next-generation sequencing (mNGS) of cerebrospinal fluid (CSF), supported by a statistical framework generated from mNGS sequencing of non-infectious patients and environmental controls, identified a pathogen.

**Design:** Case series. Using mNGS data from the CSF of 94 non-infectious neuroinflammatory cases and 24 water and reagent controls, we developed and implemented a weighted scoring metric based on z-scores at the species and genus level for both nucleotide and protein databases to prioritize and rank mNGS results. We performed mNGS on total RNA extracted from CSF of patients with subacute or chronic meningitis and highlight seven cases representing a diverse array of pathogens.

**Setting:** A multi-center study of mNGS pathogen discovery in patients with suspected neuroinflammatory conditions.

**Participants:** Patients with diagnostically challenging subacute or chronic meningitis enrolled in a research study of mNGS performed on CSF.

**Intervention:** mNGS was performed on total RNA extracted from CSF (0.25-0.5 mL). A weighted z-score was used to filter out environmental contaminants and facilitate efficient data triage and analysis.

**Main Outcomes:** 1) Pathogens identified by mNGS and 2) ability of a statistical model to prioritize, rank, and simplify mNGS results.

**Results:** mNGS identified parasitic worms, fungi and viruses in seven subjects: *Taenia solium* (n=2), *Cryptococcus neoformans*, human immunodeficiency virus-1, *Aspergillus oryzae*, *Histoplasma capsulatum*, and *Candida dubliniensis*. Evaluating mNGS data with a weighted z-score based scoring algorithm effectively separated bona fide pathogen sequences from spurious environmental sequences.

**Conclusions and Relevance:** mNGS of CSF identified a diversity of microbial pathogens in patients with diagnostically challenging subacute or chronic meningitis, including a case of subarachnoid neurocysticercosis that defied diagnosis for one year, the first case of CNS vasculitis caused by *Aspergillus oryzae*, and the fourth reported case of *Candida dubliniensis* meningitis. Filtering metagenomic data with a scoring algorithm greatly clarified data interpretation and highlights the difficulties attributing biological significance to organisms that may be present in control samples used for metagenomic sequencing studies.

**Key Points:** **Question:** How can metagenomic next-generation sequencing of cerebrospinal fluid be leveraged to aid in the diagnosis of patients with subacute or chronic meningitis?

**Findings:** Metagenomic next-generation sequencing identified parasitic worms, fungi and viruses in a case series of seven subjects. A database of water-only and healthy patient controls enabled application of a z-score based scoring algorithm to effectively separate bona fide pathogen sequences from spurious environmental sequences.

**Meaning:** Our scoring algorithm greatly simplified data interpretation in a series of patients with a wide range of challenging infectious causes of subacute or chronic meningitis identified by metagenomic next-generation sequencing.

## Introduction

Subacute and chronic meningitis is diagnostically challenging given the wide range of potential infectious, autoimmune, neoplastic, paraneoplastic, parameningeal and toxic causes.^1,2^ Securing a final diagnosis can require weeks or months of testing, and in many cases is never achieved, necessitating empiric treatment approaches that may be ineffective or even harmful.

Unlike traditional testing for a specific microbe or category of infection, metagenomic next-generation sequencing (mNGS) of cerebrospinal fluid (CSF) or brain tissue screens for all potential CNS pathogens, and can identify novel or unexpected infections in people suffering from unexplained meningitis and/or encephalitis.^3-10^ Multiple computational algorithms and pipelines have been developed to rapidly identify microbial sequences in mNGS datasets.^11-13^ However, the unbiased nature of mNGS data requires careful analysis to determine which, if any, of the identified microbes represent a true pathogen versus an environmental contaminant. Failure to make this distinction has resulted in disease associations with organisms later determined to be laboratory contaminants.^14-16^

Here, we developed a straightforward statistical approach leveraging an extensive mNGS database of water-only controls (n=24) and surplus CSF samples (n=94) obtained from patients with clinically adjudicated non-infectious neurologic diagnoses ranging from autoimmune, neoplastic, structural and neurodegenerative etiologies (“control cohort”). These data allow one to quantify how unexpected it is to identify a particular microbe at a given level of abundance in a patient sample by comparison to its mean level of abundance across the control cohort. Here we report the utility of this statistical framework for identifying microbial pathogens in seven challenging cases of subacute or chronic meningitis, as well as for analyzing publicly available data from recent mNGS infectious diagnostic and brain microbiota studies.^17-19^

## METHODS

Participants were recruited between September 2013 and March 2017 as part of a larger study applying mNGS to biological samples from patients with suspected neuroinflammatory disease. The seven participants had subacute or chronic leptomeningitis with or without encephalitis. An etiologic diagnosis was not known by the researchers at the time of study enrollment. If an infection was made by traditional means before mNGS testing was complete (participants 3, 5 and 6), the researchers performing mNGS remained blinded to the diagnosis. The cases were referred from the University of California, San Francisco (UCSF) Medical Center (n=2), Zuckerberg San Francisco General Hospital (n=2), Cleveland Clinic, University of Washington and Kaiser Permanente. The study protocol was approved by the UCSF Institutional Review Board (IRB). Participants or their surrogates provided written informed consent. Treating physicians were informed about research-based mNGS results under an IRB-approved reporting protocol. If mNGS identified a possible infection, treatment was altered after confirmatory clinical evaluation and testing.

### mNGS Protocol

mNGS was performed on total RNA extracted from surplus CSF (250-500 µL). In addition to CSF, one subject also had mNGS performed on total RNA extracted from <50 mg of snap frozen, surplus tissue from a lumbar meningeal biopsy. Samples were processed for mNGS as previously described.^5,7^ The non-human sequence reads have from each sample have been deposited at the National Center for Biotechnology Information (NCBI) Sequence Read Archive, BioProject (PRJNA338853).

### Bioinformatics

Paired-end 125-150 base pair (bp) sequences were analyzed using a previously described rapid computational pathogen detection pipeline (Figure 1).^5,7^ Unique, non-human sequences were assigned to microbial taxonomic identifiers (taxids) based on nucleotide (nt) and non-redundant (nr) protein alignments. To distinguish putative pathogens from contaminating microbial sequences derived from skin, collection tubes, lab reagents, or the environment, a composite background model of metagenomic data was employed. This model incorporated 24 water controls and 94 CSF samples from patients with non-infectious diagnoses, including 21 chronic meningitis cases with the following diagnoses: neurosarcoidosis (n=7), CNS vasculitis (n=5), autoimmune encephalitis, parameningeal dermoid cyst, leptomeningeal carcinomatosis, CNS lymphoma, melanoma, Susac syndrome, and a meningitis syndrome responsive to immunosuppression. Data were normalized to unique reads mapped per million input reads (rpM) for each microbe at the species and genus level. Using this background dataset as the expected mean rpM for a given taxid, standard Z-scores were calculated for each genus (gs) and species (sp) in each sample based on the results from both the nt and nr database searches. Thus, there are four z-scores reported for each sample: spZnt, gsZnt, spZnr and gsZnr. To prioritize reporting of the most unique (i.e., unexpected) taxa in each sample, the significance of each microbial species was mapped to a single value with the following empirically-derived formula:

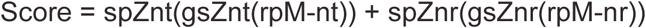

**Figure 1.**
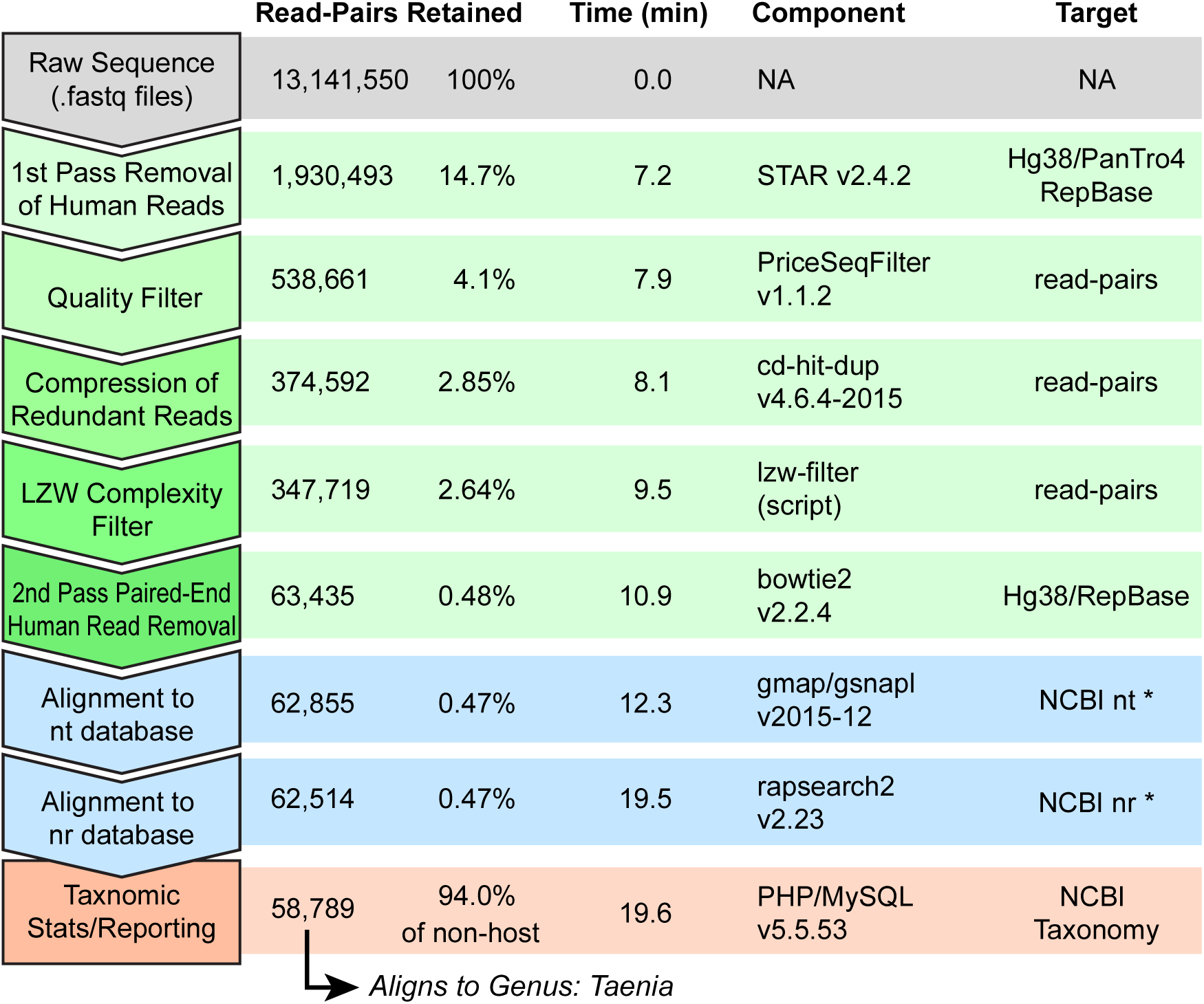
Pathogen Detection Pipeline: Parameters for Participant 1. For detection of possible pathogen-derived sequences, a rapid pipeline was implemented as diagrammed. The CSF sample from participant 1 yielded 13,141,550 million read pairs, which were then subjected to human removal (STAR v2.4.2)^24^, quality control filtering (PriceSeqFilter v1.1.2)^20^, compression of duplicate reads (cd-hit-dup v4.6.4-2015)^25^, removal of low-complexity sequences by filtering for high LZW^26^ compression ratios, a second round of human read removal (bowtie2 v2.2.4)^27^, alignment to NCBI’s nt database (gmap/gsnapl v2015-12)^28^, alignment to NCBI’s nr protein database (rapsearch2 v2.23)^29^, and finally statistical calculation and taxonomy reporting via a webpage was implemented with PHP/MySQL v5.5.53. The entire pipeline was completed in 19.6 minutes on a single high-end server (32 core, 3.2Ghz Xeon E5-2667v3, 768Gb RAM). Abbreviations: STAR, Spliced Transcripts Alignment to a Reference; LZW, Lempel-Ziv-Welch; National Center for Biotechnology Information, NCBI; nucleotide, nt; non-redundant, nr.

Here, the rpM are scaled by both the z-score for the species and the genus. If both z-scores are negative, the product remains negative. The maximum z-score is arbitrarily capped at 100. This product is calculated for alignments to both the nt and nr databases and summed. The top-ranked taxa were considered with respect to the clinical context of the patient. Microbes with known CNS pathogenicity that could cause a clinical phenotype concordant with the clinical presentation were considered potential pathogens and were confirmed by standard microbiologic assays, as described in the brief case histories below.

## RESULTS

The seven study subjects ranged in age from 10 to 55 years old, and three (43%) were female. Additional clinical details are listed in Table 1. In each case, the causative pathogen was identified within the top two scoring microbes identified by our algorithm (Figure 2).

**Table 1.** Clinical Summary. Abbreviation: lumbar puncture, LP.

| Participant | Final Diagnosis | Age | Sex | Clinical Presentation | Disease Duration at Time of Diagnostic LP (months) | Length of Follow-Up (months) |
| --- | --- | --- | --- | --- | --- | --- |
| 1 | *Taenia solium* | 28 | M | Headache and diplopia | 11 | 4 |
| 2 | *Taenia solium* | 34 | F | Headache, unilateral facial numbness and tinnitus, recurrent loss of consciousness | 9 | 21 |
| 3 | *Cryptococcus neoformans* | 52 | M | Seizures, coma | 3 | 19 |
| 4 | HIV-1 | 55 | M | Dementia, cough, night sweats, weight loss | 0.5 | 7 |
| 5 | *Aspergillus oryzae* | 32 | M | Headache, dizziness, diplopia, unilateral facial weakness and numbness | 7 | 1 |
| 6 | *Histoplasma capsulatum* | 10 | F | Back pain followed by unilateral facial droop, headache, and neck stiffness | 6 | 3.5 |
| 7 | *Candida dubliniensis* | 26 | F | Low back pain followed by saddle anesthesia and unilateral foot drop | 20 | 1 |

**Figure 2.**
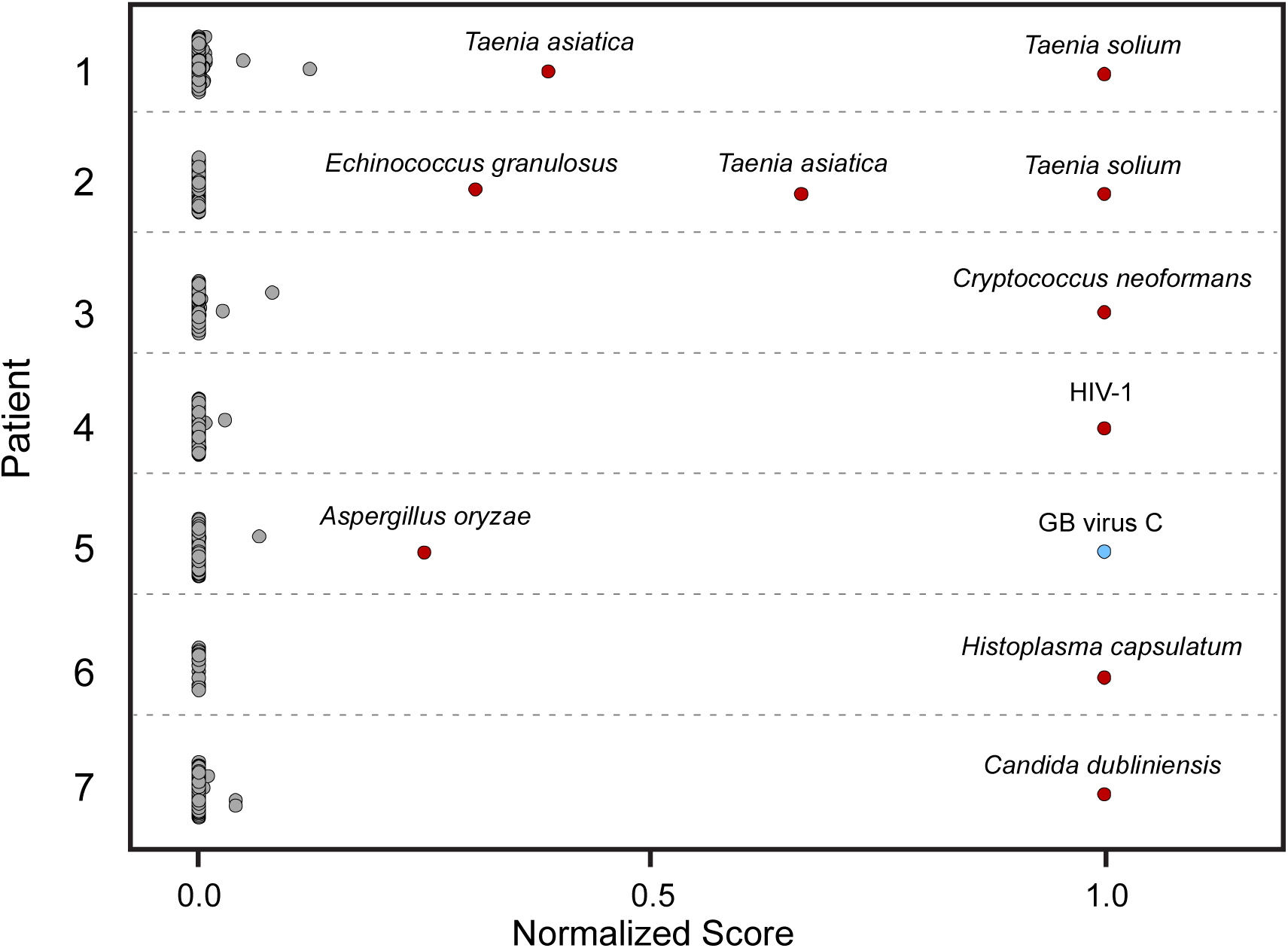
Statistical Scoring Ranking Results. Strip plot of normalized species significance scores for microbial taxa (dots) in each patient sample (row). In 6/7 samples, the neurologic infection (red dots) is ranked as the most significant by our approach. In the case of participant 5, *Aspergillus oryzae* is ranked second behind GB virus C, a likely concurrent infection unrelated to the clinical presentation.

### Case Descriptions

#### Taenia solium

Participant 1 was a 29 year-old man from Nicaragua with headache and diplopia. CSF examination revealed an opening pressure of >50 cm, 66 WBC/mm^3^ (89% lymphocytes, 4% neutrophils, 4% monocytes and 3% eosinophils), 1 RBC/mm^3^, total protein 43 mg/dL (normal range 15-50 mg/dL), and glucose 27 mg/dL (normal range 40-70 mg/dL).

Contrast-enhanced brain magnetic resonance imaging (MRI) revealed enhancement of the basilar meninges and several cranial nerves (Figure 3A). Although his serum cysticercosis antibody was positive, there were no cysts or calcifications on brain MRI. The low CSF glucose, basilar meningitis, positive tuberculin skin test, positive tuberculosis interferon-gamma release assay, and his high-risk country of origin prompted empiric treatment for *Mycobacterium tuberculosis* (TB) meningitis. He clinically improved over the next three weeks but then worsened again, requiring multiple lumbar punctures (LPs), MRIs and hospitalizations over the next year. Subsequent CSF samples showed worsening lymphocytic pleocytosis, persistently low glucose and elevated protein. Neuroimaging demonstrated persistent basilar meningitis and development of communicating hydrocephalus, again without cysts. His incomplete clinical response was initially attributed to treatment noncompliance, but after he worsened despite directly observed TB therapy, multi-drug resistant TB was suspected. After his fourth clinical decline (including discussion of ventriculoperitoneal shunt placement for worsening hydrocephalus) he was readmitted to the hospital for a new diagnostic work-up. In addition, his empiric therapy was broadened to include anti-helminthic treatment. CSF was submitted for research-based mNGS. The mNGS data demonstrated no sequences aligning to mycobacterial species. However, 58,789 unique, non-human read pairs aligned to the genus Taenia (Table 2), with the vast majority specifically mapping to the *Taenia solium* genome (Figure 2). A CSF specimen that had been sent concurrently for fungal 18s rRNA polymerase chain reaction (PCR) unexpectedly amplified the *Taenia solium* 18s rRNA gene, and a CSF cysticercosis IgG antibody was positive. After eight months of anti-helminthic treatment, he clinically improved to baseline except for a residual action tremor.

**Figure 3.**
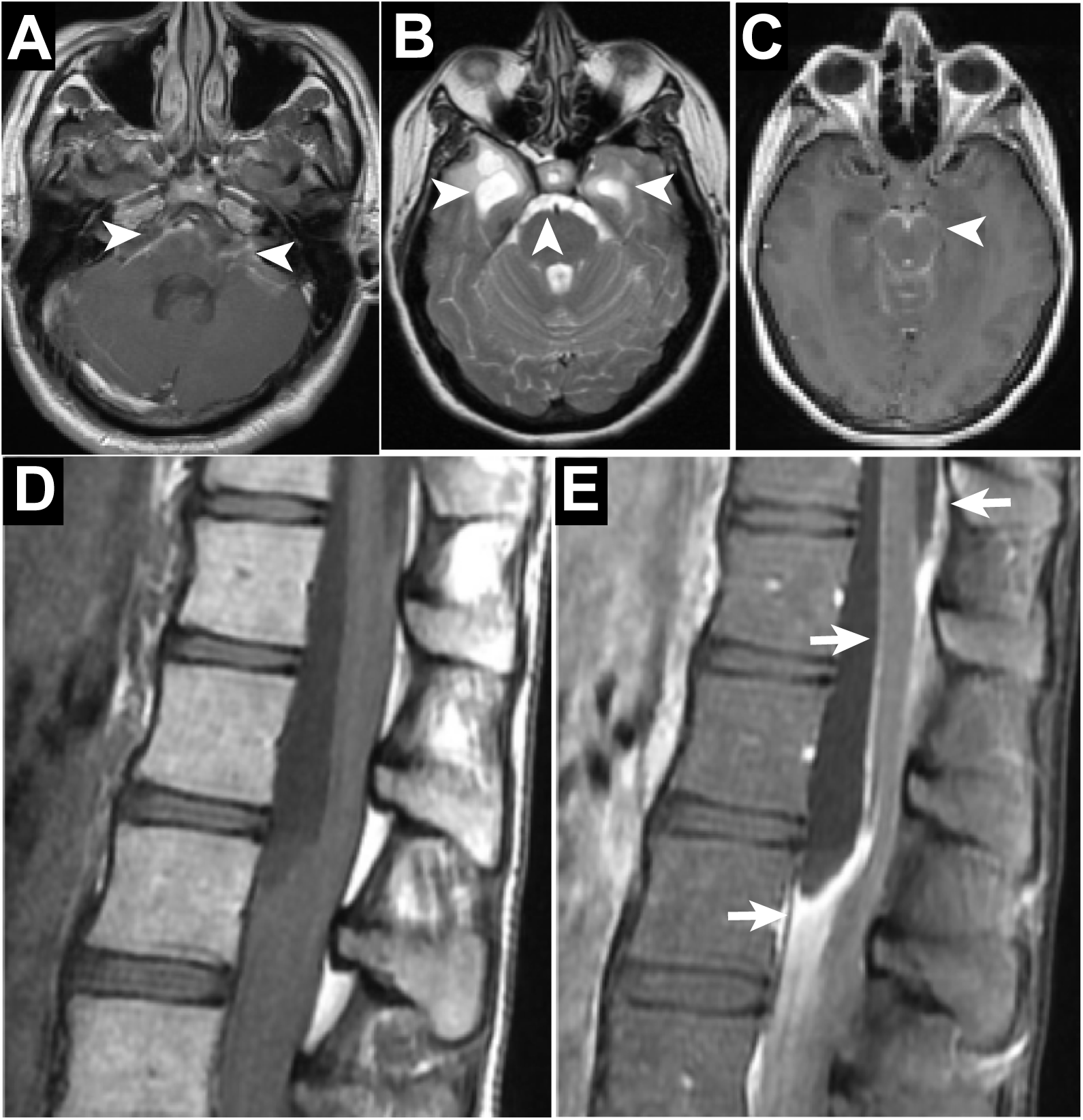
Neuroimaging. A) A 28 year-old man with neurocysticercosis identified by mNGS demonstrating basilar meningitis with contrast enhancement (arrows) on an axial post-contrast T1-weighted brain MRI, B) A 34 year-old woman with neurocysticercosis identified by mNGS demonstrating right anterior temporal lobe and pre-pontine cysts (arrows) on axial T2-weighted brain MRI, C-E) A 26 year-old woman with *Candida dubliniensis* meningitis identified by mNGS demonstrating basilar meningitis with contrast enhancement (C) (arrow), and a loculated, rim-enhancing collection extending from the top of the lumbar spinal cord anteriorly compressing the conus medullaris against the posterior wall on a sagittal pre- (D) and post- (E) contrast T1-weighted lumbar spine MRI (arrows). Abbreviation: metagenomic next-generation sequencing, mNGS; magnetic resonance imaging, MRI.

**Table 2.** Metagenomic Sequencing Summary.

| Participant | Pathogen | Total Paired-End Read Pairs | Unique non-human, non-redundant read-pairs | Unique pathogen read-pairs (% non-human read pairs) |
| --- | --- | --- | --- | --- |
| 1 | *Taenia solium* | 13,141,550 | 63,435 | 58,789 (92.7) |
| 2 | *Taenia solium* | 17,712,171 | 3,732 | 569 (15.2) |
| 3 | *Cryptococcus neoformans* | 11,121,312 | 1678 | 839 (50.0) |
| 4 | HIV-1 | 8,529,421 | 261,847 | 136,000 (51.9) |
| 5 | *Aspergillus oryzae* | 14,698,597 | 2,753 | 857 (31.1) |
| 6 | *Histoplasma capsulatum* | 13,385,787 | 9,406 | 33 (0.35) |
| 7 | *Candida dubliniensis* | 14,726,483 | 7,636 | 68 (0.89) |

Participant 2 was a 34 year-old woman who had immigrated to the United States from El Salvador 13 years prior. She presented with nine months of right-sided headache, left facial numbness, right pulsatile tinnitus and recurrent loss of consciousness. Brain MRI showed hydrocephalus and right anterior temporal lobe and pre-pontine cysts (Figure 3B). CSF examination revealed an opening pressure of 36 cm, 115 WBC/mm^3^ (1% neutrophils, 31% monocytes, 46% lymphocytes, 21% plasmacytoid lymphocytes and 1% eosinophils), 2 RBC/mm^3^, glucose <10 mg/dL and total protein 89 mg/dL. After her CSF and serum cysticercosis IgG antibody returned positive, she was treated with albendazole and prednisone for more than one month. However, she developed worsening neck pain, and a repeat CSF exam showed elevated intracranial pressure, pleocytosis with a new eosinophilia, undetectable glucose and elevated protein, raising concern for an alternative diagnosis. Her CSF mNGS data contained 569 read-pairs (Table 2) aligning to the genus Taenia (Figure 2). She was treated with dual anti-helminthic therapy with adjunctive glucocorticoids and had an excellent clinical response.

#### Cryptococcus neoformans

Participant 3 was a 52 year-old man with a history of HIV-1 infection diagnosed in 2013 (viral load detectable but too low to quantify; CD4 count 20 cells/mm^3^). He also had a history of injection drug use (IDU), hepatitis C virus infection, *Staphylococcus aureus* endocarditis, syphilis and *Cryptococcus neoformans* meningitis, the latter treated five months prior. He presented with agitation, confusion and ataxia. Clinical suspicion was high for recurrent cryptococcal meningitis, but because of his prior *S. aureus* bacteremia, syphilis and immunosuppressed state, the differential diagnosis remained broad. mNGS on CSF identified 839 unique read pairs (Table 2) that aligned to the genus Cryptococcus, essentially all of which aligned to *Cryptococcus neoformans*. Serum cryptococcal antigen was also positive at a titer of > 1:1280. Except for hepatitis C virus, no other pathogens were identified via CSF mNGS (Figure 2) or by standard diagnostic assays. He clinically improved on additional anti-cryptococcal therapy.

### Recurrent Encephalopathy in the Context of Known HIV-1 Infection

Participant 4 was a 55 year-old man with treatment-naïve HIV-1 infection diagnosed eight years prior. He presented to the Emergency Department with two weeks of confusion, decreased verbal output, as well as several months of cough, intermittent night sweats and weight loss; his CD4 count was 6 cells/mm^3^. Brain MRI showed bilateral, confluent and non-enhancing white matter T2 hyperintensities. CSF examination showed 0 WBC/mm^3^, 0 RBC/mm^3^, glucose 34 mg/dL, and total protein 93 mg/dL. CSF mNGS testing revealed more than 136,000 unique read pairs aligning to HIV-1 (Table 2, Figure 2). The full-length HIV-1 genome was assembled using paired read iterative contig extension (PRICE) v1.1.2.^20^ The mean total read coverage across the HIV-1 genome was 3,991. No reads to JC virus, herpesviruses or fungal pathogens were observed. His clinical course was consistent with HIV-1 dementia, and no other opportunistic CNS infection was identified despite extensive testing.

#### Aspergillus oryzae

Participant 5 was a 32 year-old man who presented to the Emergency Department with seven months of episodic dizziness, diplopia, headache, left facial numbness and weakness. He had a history of IDU and hepatitis C virus infection. HIV-1 testing was negative. Brain MRI revealed contrast enhancement in the left pons, middle cerebellar peduncle, and right posterior aspect of the pituitary infundibulum. A computed tomography (CT) angiogram revealed multiple areas of focal stenosis in the posterior circulation. CSF examination showed 95 WBC/mm^3^ (77% lymphocytes, 13% neutrophils, 6% monocytes, 4% reactive lymphocytes), 0 RBC/mm^3^, total protein of 61 mg/dL, glucose of 45 mg/dL. Bacterial and fungal cultures were negative for CSF and blood. He was treated for a suspected autoimmune process with glucocorticoids. Two weeks later the patient’s symptoms worsened. Repeat brain MRI revealed a new punctate infarct in the right thalamus. Repeat CSF examination showed worsening pleocytosis with a neutrophilic predominance. Bacterial cultures were negative. CSF Aspergillus galactomannan returned at 11.26 (positive > 0.5). A third CSF exam showed an even greater pleocytosis of 343 WBC/mm^3^ (63% lymphocytes, 20% neutrophils, 15% monocytes, 2% reactive lymphocytes). CSF Aspergillus galactomannan was again elevated at 7.23. CSF mNGS returned 857 read-pairs mapping to Aspergillus spp. (Table 2) with the majority of the reads mapping to *Aspergillus oryzae*. CSF 18s rRNA PCR also returned positive for Aspergillus spp., and the CSF 1,3-ß-D-glucan was elevated. No cause of immunocompromise was identified. Treatment with oral voriconazole was initiated, and his hydrocephalus was treated with ventriculoperitoneal shunting. He was then lost to follow-up.

#### Histoplasma capsulatum

Participant 6 was a 10 year-old girl with an early childhood spent in Indiana and Ohio, who presented in Seattle with six months of back pain and 10 days of progressive left facial droop, headache, neck pain, and vomiting. She had severe meningismus on exam. Brain and spinal MRI showed diffuse leptomeningeal enhancement. CSF examination revealed 68 WBC/mm^3^ (33% neutrophils and >30% monocytes), 98 RBC/mm^3^, glucose <20 mg/dL, and total protein 292 mg/dL. Empiric therapy was initiated for bacterial meningitis, herpes simplex virus, and endemic mycoses. Despite this, she developed respiratory failure and paraplegia. Repeat neuroimaging demonstrated new pontine infarcts and non-enhancing, longitudinal T2 hyperintensities throughout the cervicothoracic cord. Her therapy was broadened to cover TB meningitis. Ultimately, CSF fungal culture and 18s rRNA PCR revealed *Histoplasma capsulatum*. mNGS showed 33 read-pairs mapping to *Histoplasma capsulatum* (Table 2, Figure 2). After two months of treatment, she could ambulate but continues to have profound left hearing loss, facial weakness, and a neurogenic bladder.

#### Candida dubliniensis

Participant 7 was a 26 year-old woman with an undisclosed history of IDU who presented with one year of atraumatic lower back pain followed by subacute development of saddle anesthesia and left foot drop. On MRI, she had a loculated, rim-enhancing collection extending from the top of the lumbar spine anteriorly compressing the conus medullaris against the posterior wall, in addition to diffuse leptomeningitis involving the entire spinal cord and brainstem (Figure 3C-E). Cisternal CSF showed 126 WBC/mm^3^ (67% neutrophils, 22% lymphocytes and 11% monocytes), 4 RBC/mm^3^, total protein of 105 mg/dL, and a glucose of 40 mg/dL. Extensive infectious disease diagnostic studies were unrevealing, and 11 weeks later the patient underwent lumbar meningeal biopsy. The pathology revealed non-inflammatory, dense fibrous tissue, and no microbes were identified. CSF and tissue biopsy 18s rRNA and 16s rRNA PCRs were negative. In addition, mNGS of biopsy tissue did not reveal an infection. Three months later the patient became wheelchair-bound. Repeat cisternal CSF showed 700 WBC/mm^3^ (81% neutrophils, 16% lymphocytes, 2% monocytes and 1% eosinophils), 2 RBC/mm^3^, total protein of 131 mg/dL and a glucose of 44 mg/dL. CSF mNGS revealed 68 read-pairs mapping to Candida spp. (Table 2) with 61 of the 68 pairs mapping to *Candida dubliniensis* with 99-100% identity. A CSF 1,3-ß-D-glucan assay was 211 pg/mL (<80 pg/mL), whereas the serum 1,3-ß-D-glucan assay had been repeatedly normal.

Repeat CSF 18s rRNA and 16s rRNA PCRs were negative. The patient is being treated with combination anti-fungal therapy with mild clinical improvement, normalization of her CSF profile (including a negative CSF 1,3-ß-D-glucan) and decreasing leptomeningeal enhancement on MRI. Two of the previous three reported cases of *Candida dubliniensis* meningitis were also in patients with a history of IDU.^21-23^

### Background Signature of Reagent and Environmental Contaminants

Examination of nucleotide alignments generated by non-templated water-only controls (n=24) and non-infectious CSF samples (n=94) revealed 4,400 unique bacterial, viral, and eukaryotic genera (Supplemental Data Tables). This microbial background signature was predominated (>70%) by consistent proportions of bacterial taxa, primarily the Proteobacteria and Actinobacteria classes (Supplemental Figure 1 A, B) representing common soil, skin, and environmental flora previously reported as laboratory and reagent contaminants.^16^ To determine if these common microbial contaminants may have been misclassified as pathogens in previously published studies, we examined publicly available data from two cases of meningoencephalitis for which a possible infection was identified by mNGS.^19^ In each case, neither organism (*Delftia acidovorans*, *Elizabethkingia*) was present at levels significantly greater than the mean of our background dataset of water-only and non-infectious CSF controls (Supplementary Figure 1D). We then examined data from a study aiming to characterize the “brain microbiome” and correlate brain dysbiosis to disease.^17,18^ The abundance of the purported brain microbiota reveal distributions that are well within the observed variance we observe within our set of background water-only controls (Supplementary Figure 1C). Of note, the authors of the brain microbiome study did not deep sequence water controls from which they could not generate measurable quantities of DNA after reverse transcription-PCR (RT-PCR). The presence of environmental contaminants may likely be a function of low amounts of input RNA, which is often the case with acellular CSF samples. To test this explicitly, we performed an RNA doping experiment (Supplemental Figure 2) on a water sample and an uninfected CSF sample from which there was no detectable cDNA after RT-PCR. The mNGS libraries made from the water and CSF samples had 9.4% and 7.6% unique, non-human sequences, respectively. The proportion of non-human sequences dropped dramatically after spiking in only 20 picograms of RNA of a known identity suggesting that non-human environmental sequences are particularly problematic for low input nucleic acid samples.

## Discussion

We present seven diagnostically challenging cases of subacute and chronic meningitis in which mNGS of CSF identified a pathogen, including a case of subarachnoid neurocysticercosis that defied diagnosis for one year, the first case of CNS vasculitis caused by *Aspergillus oryzae*, and the fourth reported case of *Candida dubliniensis* meningitis. A straightforward statistical model leveraging a large mNGS dataset obtained from water-only controls and patients with a variety of non-infectious neuroinflammatory syndromes correctly prioritized the pathogens.

CSF mNGS has the potential to overcome several limitations of conventional CNS infectious disease diagnostics. First, the inherent risks of brain and/or meningeal biopsy make CSF mNGS a particularly attractive option for patients with suspected CNS infection. Second, the large number of neuroinvasive pathogens that cause subacute or chronic meningitis makes it logistically challenging and cost-prohibitive to order every possible neuro-infectious diagnostic test using a candidate-based approach. Third, some assays lack sensitivity in the context of impaired immunity or acute infection (e.g., West Nile virus serology), can be slow to yield results (e.g., mycobacterial and fungal cultures) or may fail to differentiate between active infection and prior exposure (e.g., cysticercosis antibody or the interferon-gamma release assay test for *M. tuberculosis*).

The unbiased nature of mNGS makes the datasets inherently polymicrobial and complex. Thus, statistical scoring and filtering is essential to enhance the ability to discriminate between insignificant contaminants and true infectious organisms. Our algorithm correctly prioritized etiologic pathogens in these seven clinically-confirmed cases of infectious meningitis despite the fact that the pathogens ranged widely with regard to their absolute abundance (33-136,000 sequence read pairs) and the proportion of the non-human sequences (0.89-92.7%) that they comprised (Table 2).

In addition, we analyzed a recently published clinical mNGS dataset to highlight that a thoroughgoing profile of the microbes present in water-only controls and non-infectious CSF reinforces the skepticism with which the authors described a possible infection in one subject with *Delftia acidovorans* (patient 2) and in another subject with *Elizabethkingia* (patient 7) (Supplementary Figure 1D). Furthermore, such a database could help improve the accuracy of microbiome studies, especially for body sites historically considered sterile in which rigorous controls are necessary to establish that observed microbial sequences represent microbiota vs environmental contaminants (Supplementary Figure 1C).^18,19^ This problem appears to be particularly acute in samples like CSF whose sub-nanogram levels of input RNA/DNA require unbiased molecular amplification steps before enough material is available for sequencing applications.

Indeed, the addition of only 20 picograms of purified RNA to a CSF sample was sufficient to suppress the majority of non-CSF reads deriving from the water and reagents (Supplemental Figure 2). While amplifying the input signal increases the sensitivity of the assay, it also often over-represents the signature of contaminating taxa unique to a given laboratory, experimenter, or reagent lot.^14,16^. These results provide a cautionary note and underscore the need for appropriate controls to aid in interpretation.

We expect larger databases of patient mNGS results will only enhance the ability to discriminate between irrelevant sequences and legitimate pathogens and permit more rigorous and probabilistic models for pathogen ranking and reporting. We present here one empirically derived system for prioritizing results, based on the read count weighted by standard z-scores. Given the sensitivity of NGS-based approaches, we anticipate individual laboratories will need to develop their own dynamic reference datasets to control for contaminants that are relevant to the particular time, place, and manner in which the biological samples are being analyzed.

mNGS represents an increasingly rapid and comparatively low cost means of screening CSF in an unbiased fashion for a broad range of human pathogens using a single diagnostic test. With characterization of the assay’s performance characteristics in prospective, cohort studies (http://www.ciapm.org/project/precision-diagnosis-acute-infectious-diseases), mNGS may also prove to be helpful in supporting the exclusion of CNS infection when a co-infection is suspected in an immunosuppressed patient (as illustrated by our cases with *C. neoformans* and HIV-1) or when a non-infectious cause, such as an autoimmune condition, is clinically favored. On this basis, we foresee the replacement of many single-agent assays performed in reference labs with a unified mNGS approach.

## Acknowledgements

Research reported in this article was supported by the UCSF Center for Next-Gen Precision Diagnostics supported by the Sandler Foundation and William K. Bowes, Jr. Foundation (J.L.D., M.R.W., J.M.G., F.C.C., H.A.S., K.C.Z., L.M.K.); the Rachleff Foundation (M.R.W.), Chan Zuckerberg Biohub (J.L.D.); NIH National Center for Advancing Translational Sciences (award number KL2TR000143, M.R.W.); and NIH National Institutes for Neurological Disorders and Stroke (award number K08NS096117, M.R.W.). Its contents are solely the responsibility of the authors and do not necessarily represent the official views of the NIH.

The authors of this manuscript have conflicts of interest to disclose. J.L.D. and M.R.W. are Co-Investigators of the Precision Diagnosis of Acute Infectious Diseases study funded by the CA Initiative to Advance Precision Medicine cited in the Discussion section, H.A.S. is the Program Manager of the study and K.C.Z. is a Clinical Research Coordinator for the study. The other authors of this manuscript have no conflicts of interest to disclose.

We thank E. Chow and D. Bogdanoff of the UCSF Center for Advanced Technology for their expertise and assistance operating the Illumina sequencer; the Sandler, Rachleff and William K. Bowes, Jr. Foundations as well as the Chan Zuckerberg Initiative for their generous philanthropic support; and the patients and their families for their participation in this research program.

---

**Supplemental Figure 1.**
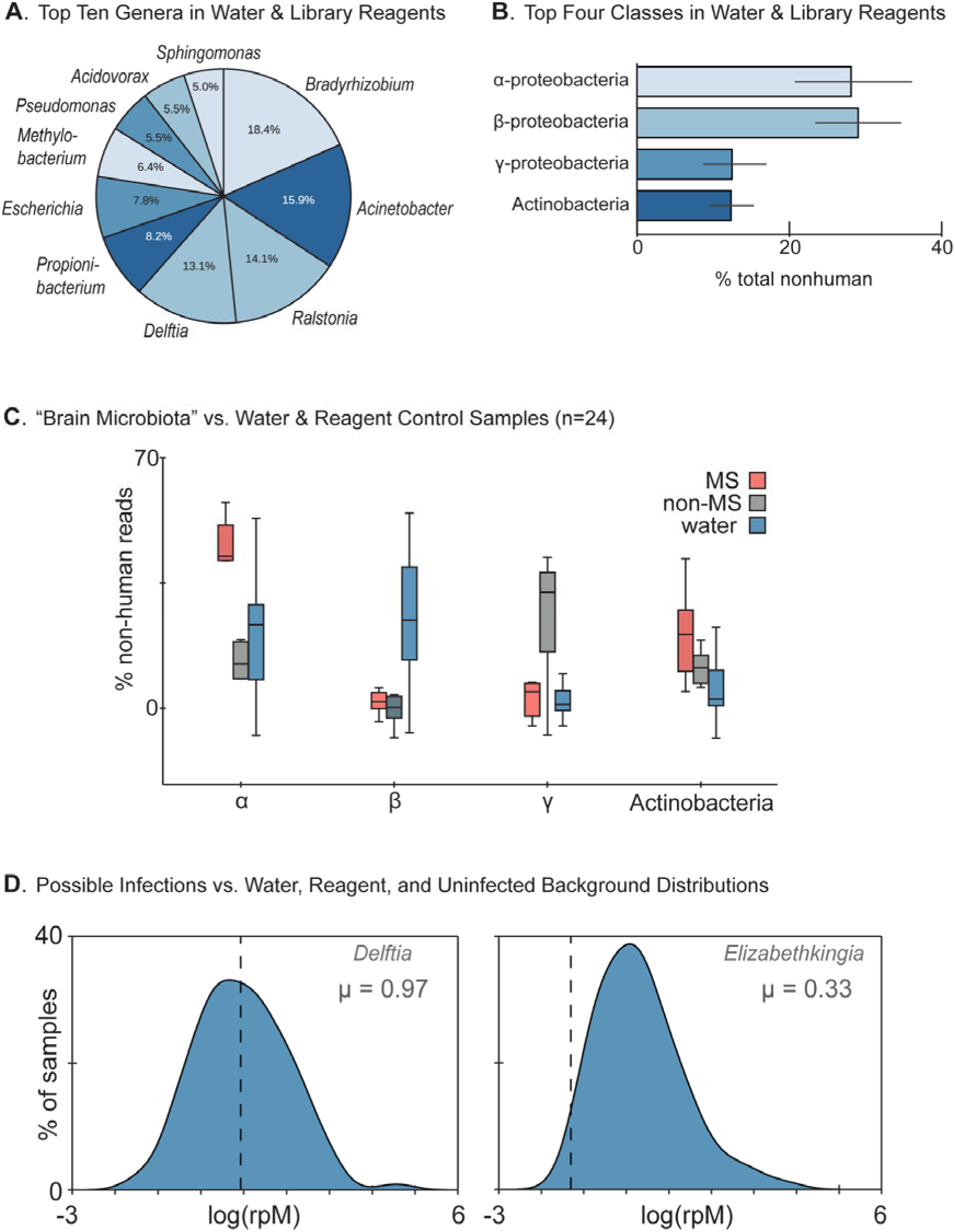
Background Signature of Reagent and Environmental Contaminants. A) 10 most abundant genera identified water-only and reagent controls (n=24). 10 bacterial taxa account for ~50% of all non-human (viral, bacterial, fungal, and selected eukaryotes) sequences in the control data. B) Organized at the class-level, four classes of bacteria (α-proteobacteria, β-proteobacteria, γ-proteobacteria, Actinobacteria) represent >80% of the sequences in the water-only and reagent controls. C) Comparison of the observed variance of the four classes of bacteria (B) to publicly available data from a recent report^18^ on microbiota in healthy and multiple sclerosis patient brain specimens suggests that the relative abundances of each taxa in the brain specimens are within the observed variance of our background dataset. D) The proportion of *Delftia acidovorans* sequences (dotted line at 0.93) in a reported case of possible CNS infection identified by mNGS^19^ is concordant with the expected mean of our control dataset (0.97). Similarly, the proportion of *Elizabethkingia* sequences (dotted line) in another possible infectious case from the same report is significantly lower than the mean abundance in our control dataset.

**Supplemental Figure 2.**
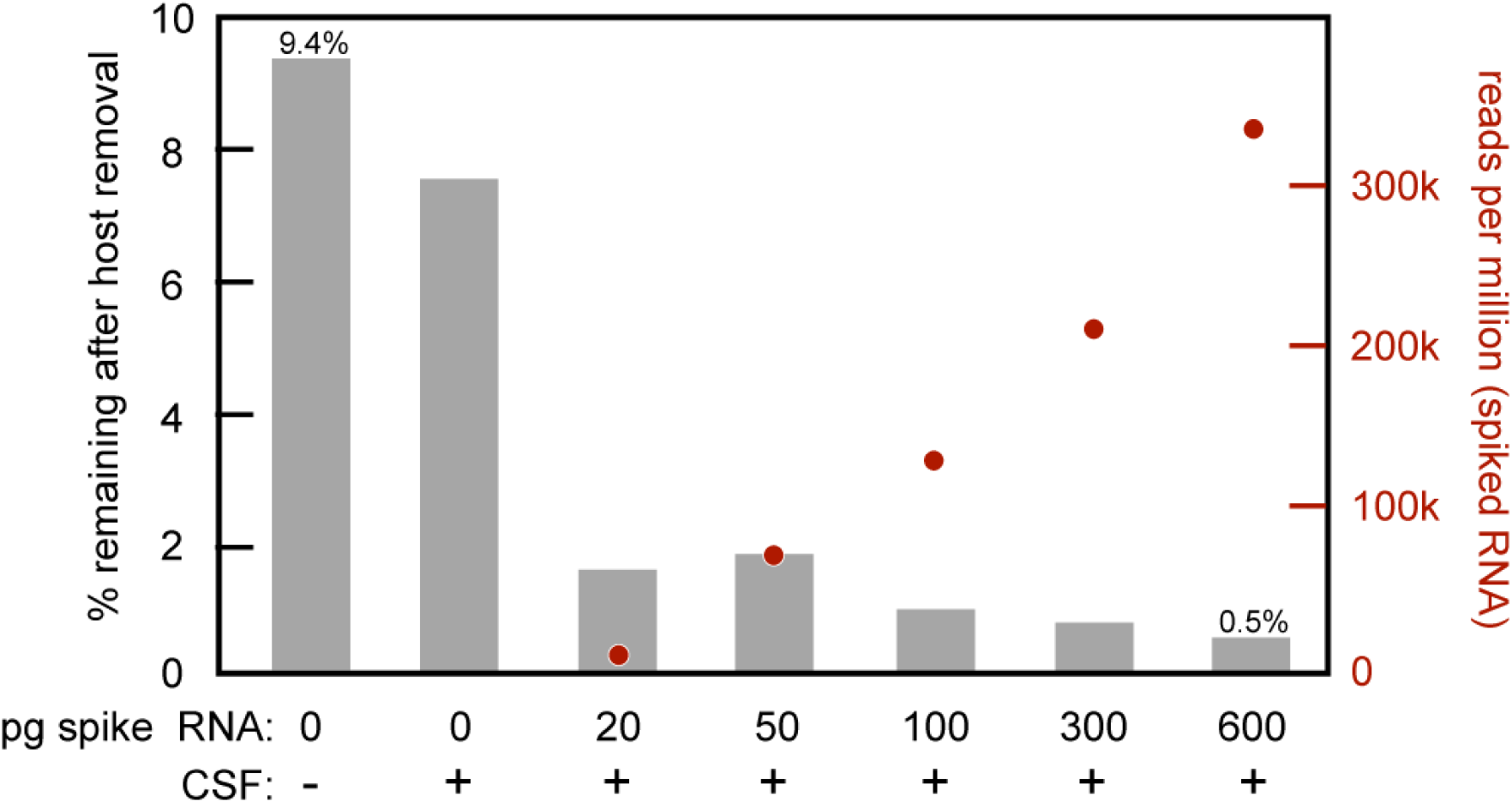
RNA Doping Experiment. Comparison of the percent non-human sequences (y-axis) found in water (column 1) and a cerebrospinal fluid (CSF) control (column 2) and the decrease in the percent non-human sequences found with increasing amounts of spiked RNA of a known identity (columns 3-7). Added RNA was generated by T7 *in vitro* transcription from a cloned luciferase reporter gene, purified, quantified, and spiked into the CSF at the indicated amounts. These data suggest that common environmental contaminants are present at low picogram quantities, and the addition of only 20 pg is sufficient to suppress the majority of reads not derived from the CSF.

## References

1. Zunt JR, Baldwin KJ. Chronic and subacute meningitis. Continuum (Minneap Minn). 2012;18(6 Infectious Disease):1290–1318.

2. Baldwin KJ, Zunt JR. Evaluation and treatment of chronic meningitis. Neurohospitalist. 2014;4(4):185–195.

3. Wilson MR, Naccache SN, Samayoa E, et al. Actionable diagnosis of neuroleptospirosis by next-generation sequencing. N Engl J Med. 2014;370(25):2408–2417.

4. Wilson MR, Shanbhag NM, Reid MJ, et al. Diagnosing Balamuthia mandrillaris Encephalitis With Metagenomic Deep Sequencing. Ann Neurol. 2015;78(5):722–730.

5. Wilson MR, Zimmermann LL, Crawford ED, et al. Acute West Nile Virus Meningoencephalitis Diagnosed Via Metagenomic Deep Sequencing of Cerebrospinal Fluid in a Renal Transplant Patient. Am J Transplant. 2016.

6. Murkey JA, Chew KW, Carlson M, et al. Hepatitis E Virus-Associated Meningoencephalitis in a Lung Transplant Recipient Diagnosed by Clinical Metagenomic Sequencing. Open Forum Infect Dis. 2017;4(3):ofx121.

7. Wilson MR, Suan D, Duggins A, et al. A novel cause of chronic viral meningoencephalitis: Cache Valley virus. Ann Neurol. 2017;82(1):105–114.

8. Naccache SN, Peggs KS, Mattes FM, et al. Diagnosis of neuroinvasive astrovirus infection in an immunocompromised adult with encephalitis by unbiased next-generation sequencing. Clin Infect Dis. 2015;60(6):919–923.

9. Palacios G, Druce J, Du L, et al. A new arenavirus in a cluster of fatal transplant-associated diseases. N Engl J Med. 2008;358(10):991–998.

10. Quan PL, Wagner TA, Briese T, et al. Astrovirus encephalitis in boy with Xlinked agammaglobulinemia. Emerg Infect Dis. 2010;16(6):918–925.

11. Flygare S, Simmon K, Miller C, et al. Taxonomer: an interactive metagenomics analysis portal for universal pathogen detection and host mRNA expression profiling. Genome Biol. 2016;17(1):111.

12. Naccache SN, Federman S, Veeraraghavan N, et al. A cloud-compatible bioinformatics pipeline for ultrarapid pathogen identification from nextgeneration sequencing of clinical samples. Genome Res. 2014;24(7):1180–1192.

13. Wood DE, Salzberg SL. Kraken: ultrafast metagenomic sequence classification using exact alignments. Genome Biol. 2014;15(3):R46.

14. Naccache SN, Greninger AL, Lee D, et al. The perils of pathogen discovery: origin of a novel parvovirus-like hybrid genome traced to nucleic acid extraction spin columns. J Virol. 2013;87(22):11966–11977.

15. Lee D, Das Gupta J, Gaughan C, et al. In-depth investigation of archival and prospectively collected samples reveals no evidence for XMRV infection in prostate cancer. PLoS One. 2012;7(9):e44954.

16. Salter SJ, Cox MJ, Turek EM, et al. Reagent and laboratory contamination can critically impact sequence-based microbiome analyses. BMC Biol. 2014;12:87.

17. Branton WG, Ellestad KK, Maingat F, et al. Brain microbial populations in HIV/AIDS: alpha-proteobacteria predominate independent of host immune status. PLoS One. 2013;8(1):e54673.

18. Branton WG, Lu JQ, Surette MG, et al. Brain microbiota disruption within inflammatory demyelinating lesions in multiple sclerosis. Sci Rep. 2016;6:37344.

19. Salzberg SL, Breitwieser FP, Kumar A, et al. Next-generation sequencing in neuropathologic diagnosis of infections of the nervous system. Neurol Neuroimmunol Neuroinflamm. 2016;3(4):e251.

20. Ruby JG, Bellare P, Derisi JL. PRICE: software for the targeted assembly of components of (Meta) genomic sequence data. G3 (Bethesda). 2013;3(5):865–880.

21. van Hal SJ, Stark D, Harkness J, Marriott D. Candida dubliniensis meningitis as delayed sequela of treated C. dubliniensis fungemia. Emerg Infect Dis. 2008;14(2):327–329.

22. Andrew NH, Ruberu RP, Gabb G. The first documented case of Candida dubliniensis leptomeningeal disease in an immunocompetent host. BMJ Case Rep. 2011;2011.

23. Yamahiro A, Lau KH, Peaper DR, Villanueva M. Meningitis Caused by Candida Dubliniensis in a Patient with Cirrhosis: A Case Report and Review of the Literature. Mycopathologia. 2016;181(7-8):589–593.

24. Dobin A, Davis CA, Schlesinger F, et al. STAR: ultrafast universal RNA-seq aligner. Bioinformatics. 2013;29(1):15–21.

25. Fu L, Niu B, Zhu Z, Wu S, Li W. CD-HIT: accelerated for clustering the nextgeneration sequencing data. Bioinformatics. 2012;28(23):3150–3152.

26. Ziv J LA. A universal algorithm for sequential data compression. IEEE Trans Inf Theory. 1977;23:337–343.

27. Langmead B, Salzberg SL. Fast gapped-read alignment with Bowtie 2. Nat Methods. 2012;9(4):357–359.

28. Wu TD, Nacu S. Fast and SNP-tolerant detection of complex variants and splicing in short reads. Bioinformatics. 2010;26(7):873–881.

29. Zhao Y, Tang H, Ye Y. RAPSearch2: a fast and memory-efficient protein similarity search tool for next-generation sequencing data. Bioinformatics. 2012;28(1):125–126.

